# Heme oxygenase-1 affects cytochrome P450 function through the formation of heteromeric complexes: Interactions between CYP1A2 and heme oxygenase-1

**DOI:** 10.1101/2020.09.14.296467

**Authors:** J. Patrick Connick, James R. Reed, George F. Cawley, Wayne L. Backes

**Affiliations:** Department of Pharmacology and Experimental Therapeutics, Louisiana State University Health Sciences Center – New Orleans, LA

**Author notes:** Corresponding author: Wayne L. Backes.

**Keywords:** Cytochrome P450, heme oxygenase, protein-protein interaction, bioluminescence resonance energy transfer (BRET), membrane protein, structure-function, electron transfer, NADPH-cytochrome P450 reductase, CYP1A2

## Abstract

Heme oxygenase 1 (HO-1) and the cytochromes P450 (P450s) are endoplasmic reticulum-bound enzymes that rely on the same protein, NADPH-cytochrome P450 reductase (POR), to provide the electrons necessary for substrate metabolism. Although the HO-1 and P450 systems are interconnected due to their common electron donor, they generally have been studied separately. As the expression of both HO-1 and P450s are affected by xenobiotic exposure, changes in HO-1 expression can potentially affect P450 function, and conversely, changes in P450 expression can influence HO-1. The goal of this study was to examine interactions between the P450 and HO-1 systems. Using bioluminescence resonance energy transfer (BRET), HO-1 formed HO-1•P450 complexes with CYP1A2, CYP1A1, and CYP2D6, but not all P450s. Studies then focused on the HO-1/CYP1A2 interaction. CYP1A2 formed a physical complex with HO-1 that was stable in the presence of POR. As expected, both HO-1 and CYP1A2 formed BRET-detectable complexes with POR. Whereas the POR•CYP1A2 complex was readily disrupted by the addition of HO-1, the POR•HO-1 complex was not significantly affected by the addition of CYP1A2. Interestingly, enzyme activities did not follow this pattern. Whereas BRET data suggested substantial inhibition of CYP1A2-mediated 7-ethoxyresorufin deethylation in the presence of HO-1, its activity was actually stimulated at subsaturating POR. In contrast, HO-1-mediated heme metabolism was inhibited at subsaturating POR. These results indicate that HO-1 and CYP1A2 form a stable complex and have mutual effects on the catalytic behavior of both proteins that cannot be explained by simple competition for POR.

## Introduction

P450 is a heme-containing, membrane-bound protein that catalyzes the oxygenation of a wide variety of exogenous and endogenous compounds (1,2). Generally, metabolism by P450 converts lipid-soluble substrates to more water-soluble products and thereby facilitates their elimination. However, in many instances, metabolism by P450 can generate reactive intermediates that lead to toxicity, mutagenesis, and/or carcinogenesis (2,3). Additionally, the standard monooxygenase reaction can be “uncoupled,” leading to generation of superoxide and H_2_O_2_ (4).

HO-1 is responsible for the first step of heme degradation, converting the substrate to biliverdin (5,6). This enzyme protects cells from oxidative damage, and is induced by agents that are related to oxidative stress, including many xenobiotics, such as aspirin, statins, niacin, etc. (7–9). HO-1 levels are also influenced by disease states including inflammation, diabetes, hepatic injury, infectious disease, and cancer (6–8). A feature common to these conditions is oxidative stress, where HO-1 induction leads to the elimination of pro-oxidants (heme) and the generation of antioxidants (e.g. bilirubin and CO).

There is evidence that HO-1 and some P450s are capable of forming homomeric complexes that can affect function. HO-1 (10,11), CYP1A2 (12), CYP2C2 (13), CYP2E1 (12,14), and CYP3A4 (15,16) were reported to form homomeric complexes, which in several cases, affected enzyme catalysis. The formation of heteromeric complexes also produce significant effects on P450 function. Such complexes have been reported for CYP1A2/CYP2B4, CYP1A2/CYP2E1, CYP2C9/CYP3A4, CYP1A1/CYP3A2, CYP2C9/CYP2D6, CYP2E1/CYP2B4, CYP2D6/CYP2E1, CYP3A4/CYP2E1, and CYP2C9/CYP3A4 (15,17–30).

Heme oxygenase and P450s do not function alone, but require an interaction with NADPH-cytochrome P450 reductase (POR) (31), which in most tissues, is found in limiting concentrations (32). Although the HO-1 and P450 systems are interconnected by their common electron donor, the systems generally have been studied separately. Whereas the existence of both physical and functional P450•P450 interactions has been well-established, potential interactions between HO-1 and the P450s have attracted relatively little attention. Still, evidence that P450s and HO-1 can affect each other’s function does exist. Anwar-Mohamed et al. (33) showed that the arsenite-induced inhibition of CYP1A1, CYP1A2, CYP3A23, and CYP3A2 in rat hepatocytes was at least partially mediated by induction of HO-1, but the mechanism for these effects was not investigated.

The known abundance of P450•P450 interactions and their observed effects on enzyme function suggest interactions between HO-1 and P450s may also influence catalytic activities. The goal of this study was to examine the potential for interactions between the HO-1 and P450 systems, focusing on CYP1A2. The results show that the presence of HO-1 has a significant impact on P450 function that cannot be explained by a simple mass-action competition between HO-1 and P450 for limiting reducing equivalents. The results also show that HO-1 and CYP1A2 form a stable complex both in reconstituted systems and in natural membranes, suggesting that the change in both CYP1A2 and HO-1 function is the result of this complex.

## Results

### Effect of HO-1 on complex formation with different P450 enzymes

With the knowledge that P450•P450 interactions have been associated with changes in protein activity, the potential for physical interactions between HO-1 and P450s was examined. In an effort to determine if interactions between different P450s and HO-1 were selective for the individual enzymes, an initial search was performed with different GFP-tagged P450s and Rluc-tagged HO-1. Because full-length HO-1 has a naturally occurring C-terminal membrane-binding region, the Rluc-HO-1 fusion protein was engineered with an N-terminal Rluc tag. Conversely, P450s and POR (having N-terminal membrane domains) were labeled on the C-terminus. This was done to minimize association of the tags with the membrane-binding regions of the proteins, allowing them to interact with the endoplasmic reticulum. HEK293T/17 cells were transfected with the proteins at a high GFP:Rluc ratio to ensure that the measured signal approximated BRETmax for each pair. Cells were incubated for 48 h post-transfection to optimize protein expression to allow for the detection of lower-affinity P450/HO-1 interactions. Under these conditions, higher BRET signals were observed where HO-1 was paired with CYP1A1, CYP2D6, and CYP1A2, with much weaker responses observed with CYP2A6 and CYP2C9 (Fig. 1). Based on these data, the HO-1/CYP1A2 reaction was examined for further study. These results suggest that interactions between HO-1 and P450s are specific for the particular P450 enzyme present.

**Figure 1.**
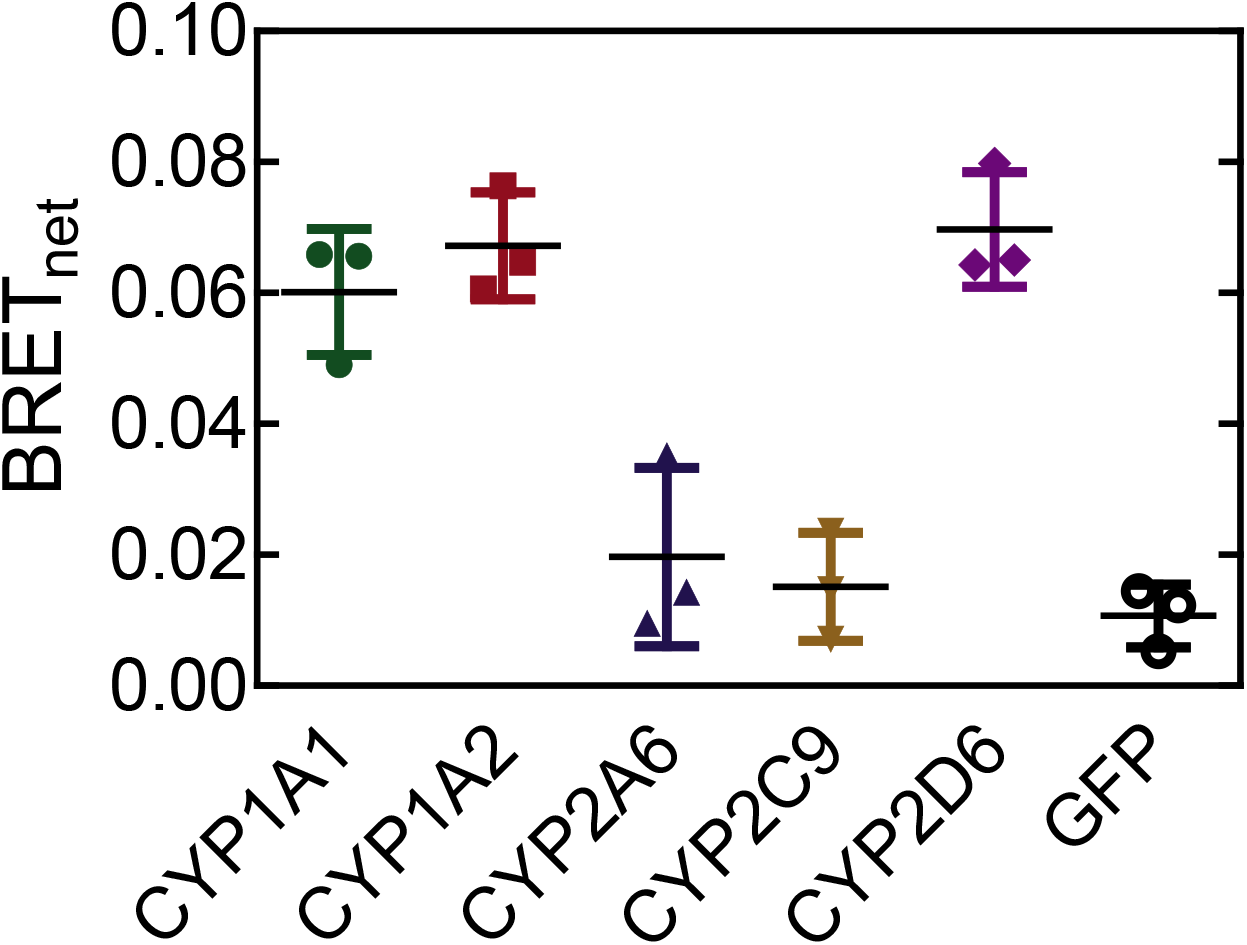
Interactions between HO-1 and different P450 enzymes. Screen of various cytochromes P450 for their potential interaction with heme oxygenase-1. HEK 293T/17 cells were transfected with vectors coding for Rluc-tagged HO-1 and GFP-tagged P450. Expression of GFP-tagged protein was between 15- and 30-fold higher than HO-1-Rluc expression for each pair in order to approach a maximal BRET response. Error bars represent the standard deviation (SD) for triplicate measurements of cells from a single transfection.

### Characterization of physical interactions among CYP1A2, HO-1, and POR

We then focused on CYP1A2 because of the high BRET signal generated when paired with HO-1 in the initial screening experiment (Fig. 1), and the known ability of CYP1A2 to form complexes with other P450s (17–21,26,34). The goal of this study was to determine whether CYP1A2 and HO-1 formed a specific BRET-detectable complex, and to see if the complex was stable in the presence of POR. This was accomplished by measuring complex formation for each of the three potential binary interactions among CYP1A2/HO-1, HO-1/POR, and CYP1A2/POR in the presence and absence of the third protein. Expression vectors for CYP1A2-GFP and Rluc-HO-1 were transfected into HEK 293T/17 cells at a range of GFP:Rluc ratios. These data follow saturation curves that were fit to a hyperbolic function. The maximum BRET value (BRETmax) correlates with the fraction of Rluc-tagged proteins interacting with GFP-tagged proteins, and serves as an indicator of protein complex formation. Adjustments to the DNA levels were made to ensure that measured differences in BRETmax could not be attributed to differences in protein expression levels. Specifically, when a difference in BRETmax was observed, total tagged protein expression for each point of the lower curve was between 1.0 and 1.2x the average total tagged protein expression of the higher curve (Fig. 2B).

**Figure 2.**
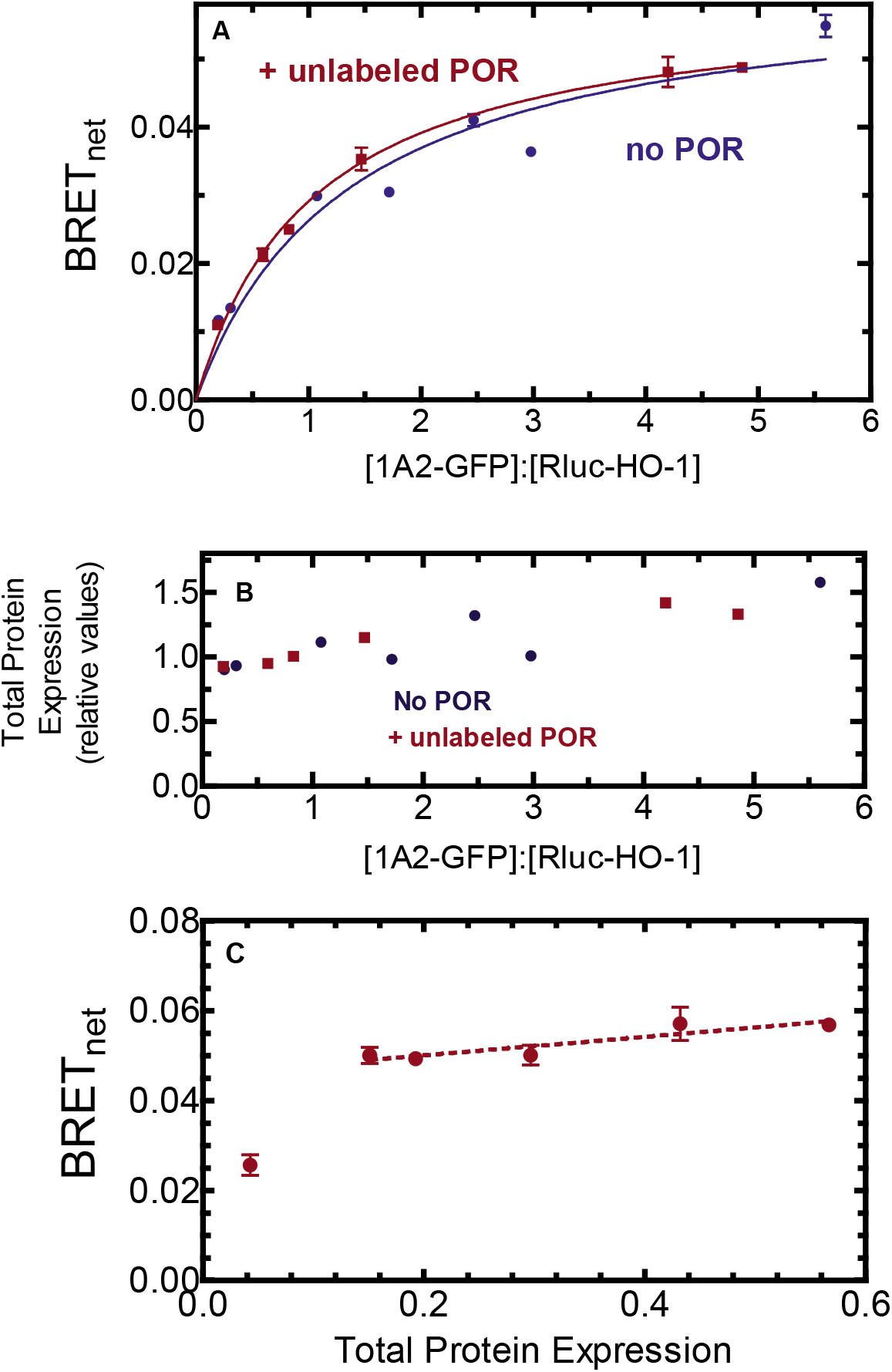
Determination of complex formation between CYP1A2 and HO-1 – Effect of POR. **A.** HEK 293T/17 cells were transfected with plasmids coding for rabbit CYP1A2-GFP, Rluc-HO-1, in the absence and presence of untagged-POR. The CYP1A2-GFP/Rluc-HO-1 BRET pair was measured 24 hours after transfection in the absence (blue), and presence of 500 ng (red) of co-transfected POR DNA. Error bars represent the represent the standard deviation (SD) of triplicate measurements of cells from a single transfection and generally do not exceed the size of the points. The experiment was performed three times with small adjustments to transfection conditions for optimization of protein expression levels. Results from each transfection were consistent. B. Total protein expression was estimated from the sum of the fluorescence of the GFP and luminescence of the Rluc tags, using a GFP-Rluc fusion protein as a standard. C. Effect of changes in protein concentration at a fixed ratio of GFP-CYP1A2/Rluc-HO-1. HEK 293T/17 cells were transfected with different amounts of DNA, while maintaining an excess of the CYP1A2-GFP tagged protein. The relative levels of GFP:Rluc protein expression were in excess of 38:1.

The first goal was to establish that HO-1 and CYP1A2 formed a stable complex, and to determine if it could be disrupted by the presence of POR. Fig. 2A shows that the CYP1A2-GFP/Rluc-HO-1 BRET pair generated a saturation curve that was consistent with the formation of a specific CYP1A2•HO-1 complex. Transfection of unlabeled POR to the HEK293T cells showed no significant disruption of this complex, suggesting that the CYP1A2•HO-1 complex was stable. Fig. 2B shows that the total amount of protein expression was similar in both the absence and presence of co-transfected, unlabeled POR.

The specificity of complex formation was also illustrated by measuring the BRET response as a function of total protein expression (Fig. 2C). If these complexes were not specific and simply the result of protein crowding, a straight line with a positive slope going through the origin would be expected (35). These results also demonstrate that over a large range of transfection levels, the maximal BRET response is relatively insensitive to changes in levels of transfected DNA. Taken together, these results indicate that HO-1 and CYP1A2 form a specific complex when co-transfected into living cells, and that the complex is stable in the presence of added POR.

Next, the effect of HO-1 on CYP1A2•POR complex formation was examined. As expected, CYP1A2 and POR formed an efficient BRET complex (Fig. 3A). Co-transfection of untagged HO-1 alongside the CYP1A2-GFP/POR-Rluc BRET pair led to a greater than 70% decrease in BRETmax. Again, levels of the proteins in the BRET pair were similar both in the absence and presence of unlabeled HO-1 (Fig. 3B), indicating that the decrease in BRET signal was not simply due to a decrease in protein expression, but due to disruption of the POR•CYP1A2 complex. The disruption of the POR•CYP1A2 complex is consistent with HO-1 preferentially binding to POR when all three proteins are present.

**Figure 3.**
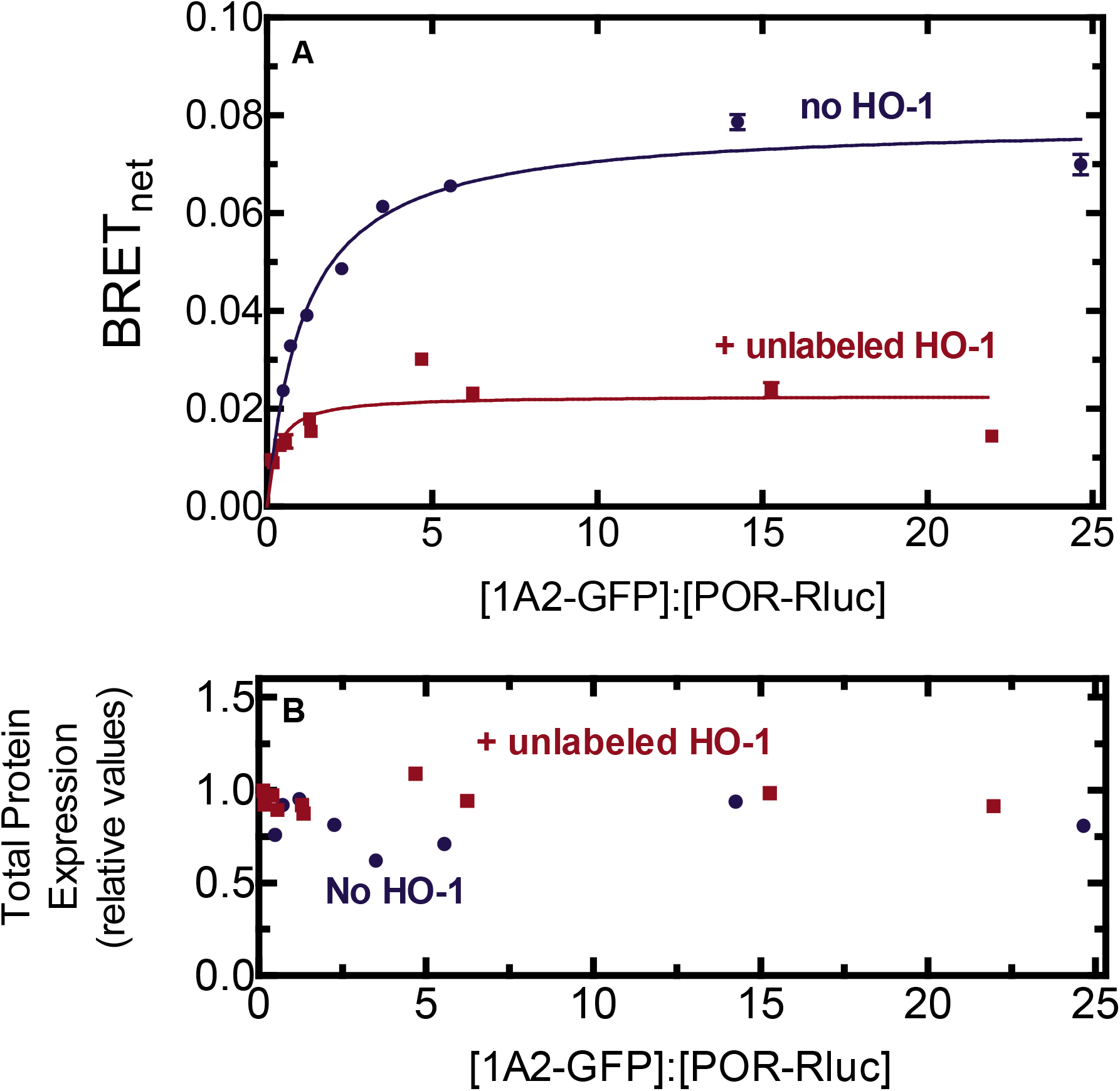
Effect of HO-1 on the interaction between CYP1A2-GFP and POR-Rluc. A. HEK 293T/17 cells were transfected with plasmids coding for rabbit CYP1A2-GFP and POR-Rluc in the absence and presence of 500 ng unlabeled HO-1. Twenty-four hours post-transfection, cells were collected, and BRET was measured. When cells were co-transfected with HO-1 DNA the maximum BRET signal generated was significantly lower (red) than that of cells transfected with the CYP1A2-GFP•POR-Rluc pair alone (blue). Data points represent triplicate measurements of cells from a single transfection; error bars represent the standard deviation (SD) and generally did not exceed the size of the points. Each experiment was repeated with small adjustments to transfection conditions for optimization of protein expression. Results were both transfections were consistent. B. Total protein expression was estimated from the sum of the fluorescence of the GFP and luminescence of the Rluc tags.

In the complementary experiment, the effect of unlabeled CYP1A2 on the POR•HO-1 BRET pair was examined. Again, the expected complex between HO-1 and POR was generated (Fig. 4). Surprisingly, co-transfection of unlabeled CYP1A2 did not significantly affect formation of the POR•HO-1 complex. Again, these results are consistent with a CYP1A2•HO-1 complex where POR selectively associates with its HO-1 moiety.

**Figure 4.**
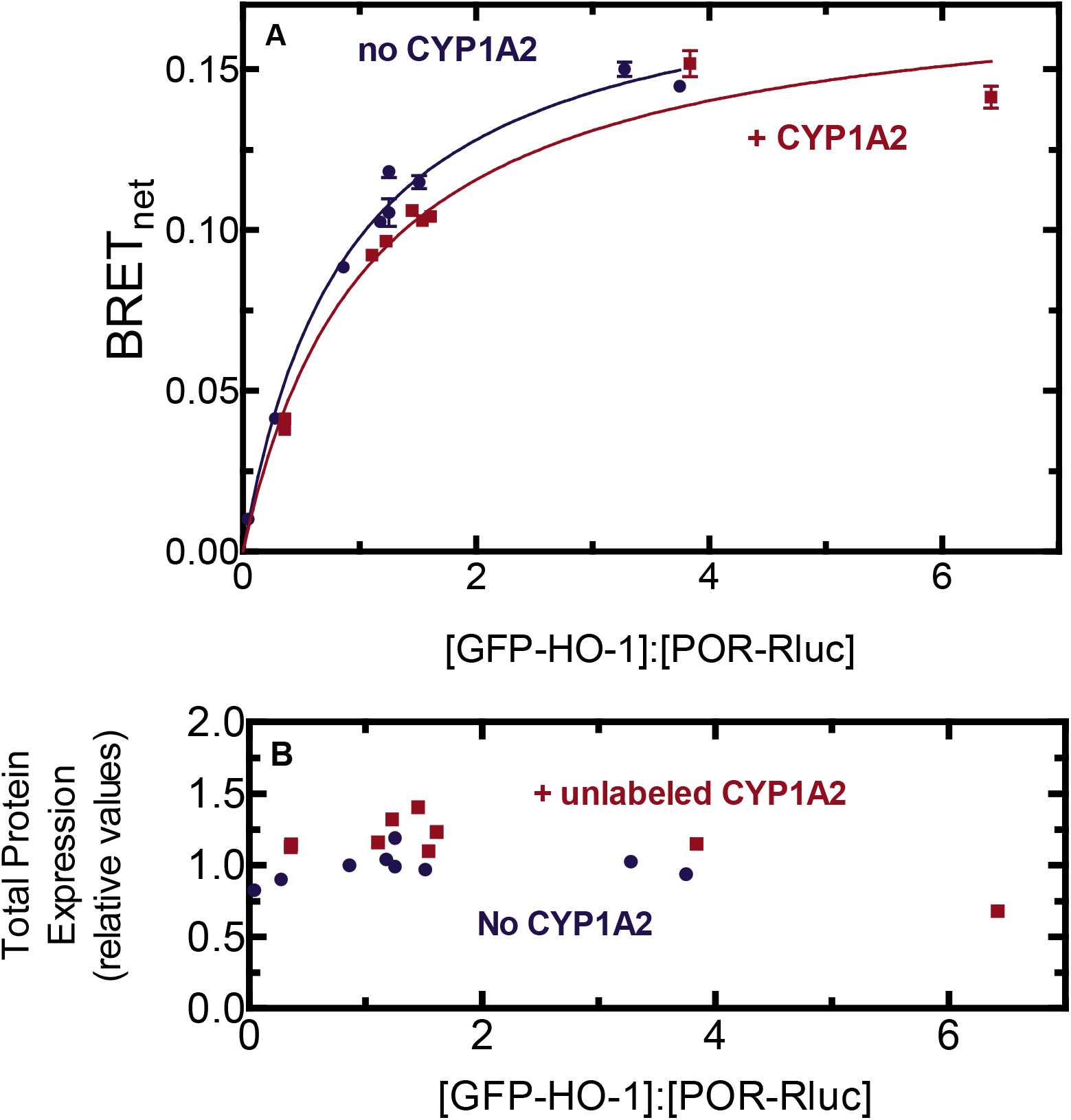
Effect of CYP1A2 on the interaction between GFP-HO-1 and POR-Rluc. A. HEK 293T/17 cells were transfected with plasmids coding for GFP-HO-1 and POR-Rluc in the absence and presence of untagged CYP1A2. BRET generated by the GFP-HO-1•POR-Rluc pair was measured 24 hours after transfection in the presence (red) and absence (blue) of 2 μg of vector coding for untagged CYP1A2. Co-transfected CYP1A2 DNA did not significantly alter the BRET signal generated by the GFP-HO-1•POR-Rluc pair. Error bars represent the SD of triplicate measurements of cells from a single transfection and generally do not exceed the size of the points. Each experiment was repeated with small adjustments to transfection conditions for optimization of protein expression, generating similar results. B. Total protein expression was estimated from the sum of the fluorescence of the GFP and luminescence of the Rluc tags.

CYP1A2 is known to form homomeric complexes in membranes, which was shown by both BRET and chemical crosslinking (12). As this homomeric BRET complex is known to be disrupted by the co-transfection of unlabeled POR, the potential of unlabeled HO-1 to disrupt the CYP1A2/CYP1A2 BRET pair was examined (Fig. 5). The results clearly show that HO-1 can disrupt the CYP1A2/CYP1A2 BRET pair.

**Figure 5.**
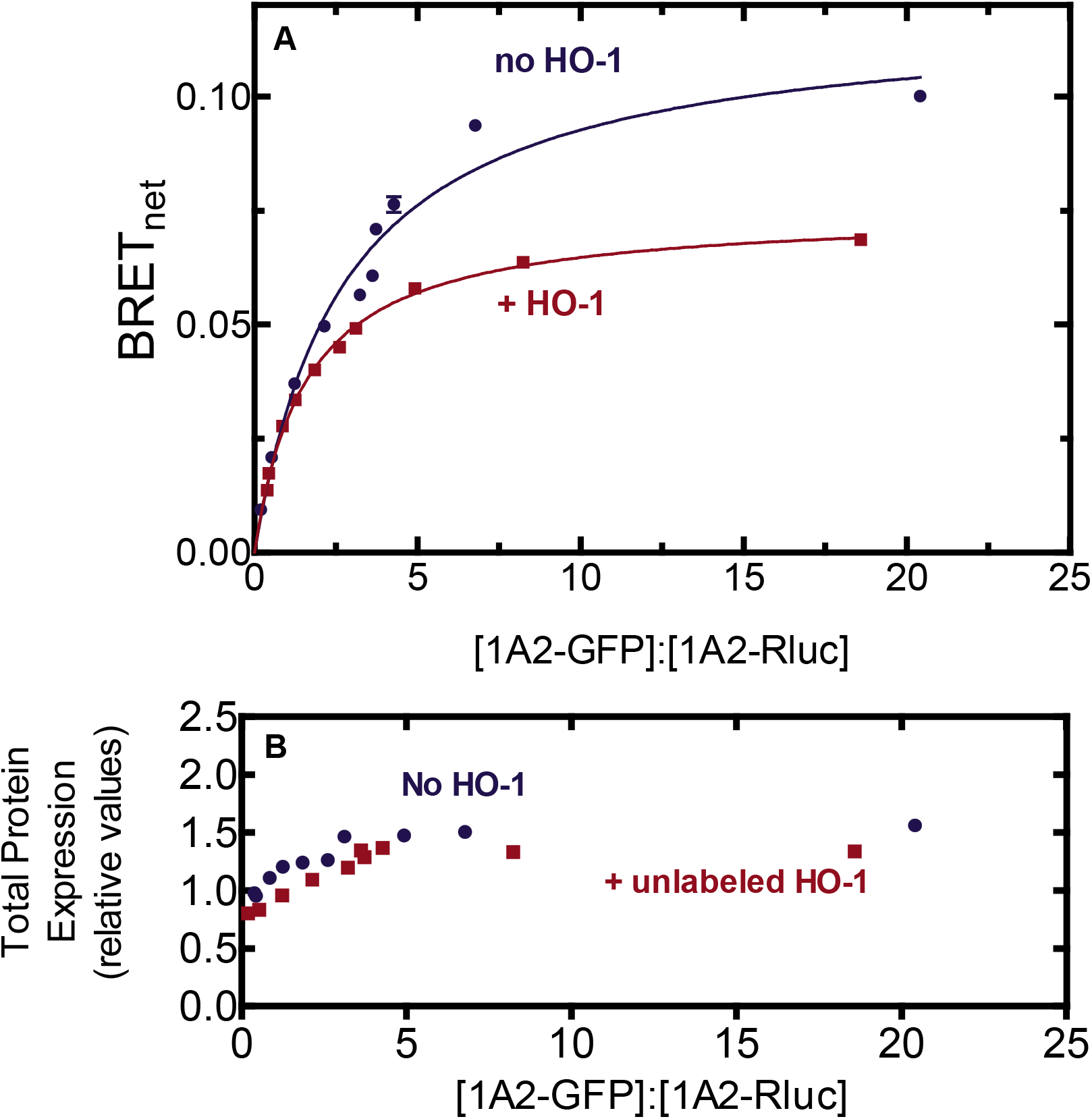
Effect of HO-1 on formation of the homomeric CYP1A2 BRET complex. A. HEK 293T/17 cells were transfected with plasmids coding for CYP1A2-GFP and CYP1A2-Rluc in the absence and presence of untagged HO-1. BRET generated by the CYP1A2-GFP•CYP1A2-Rluc pair was measured 24 hours after transfection with (red) and without (blue) 2000 ng of a vector coding for of untagged HO-1. Co-transfected HO-1 DNA led to a significant disruption of the BRET signal generated by the CYP1A2-GFP•CYP1A2-Rluc pair. Error bars represent the standard deviation (SD) of triplicate measurements of cells from a single transfection, and do not exceed the size of the data points. Each experiment was repeated with small adjustments to transfection conditions for optimization of protein expression, generating similar results. B. Total protein expression was estimated from the sum of the fluorescence of the GFP and luminescence of the Rluc tags.

### Effect of HO-1 on CYP1A2 activity

The presence of a stable CYP1A2•HO-1 complex leads to the question of what effect this complex might have on P450 and HO-1 activities. To investigate this, CYP1A2-mediated 7-ethoxyresorufin deethylation (EROD) was measured as a function of POR concentration where the proteins were reconstituted into L-*α*-dilauroyl-*sn*-glycero-3-phosphocholine (DLPC). In the absence of HO-1, CYP1A2 produced a sigmoidal response as a function of POR concentration (Fig. 6). This corroborates our previous reports and is consistent with the tendency of CYP1A2 to form a homomeric complex (Fig. 5 & (12)). Interestingly, in the presence of HO-1, EROD appeared to be stimulated at subsaturating POR, but moderately inhibited at higher POR concentrations, showing a V_max_ that was about 20% lower when HO-1 was present in the reaction.

**Figure 6.**
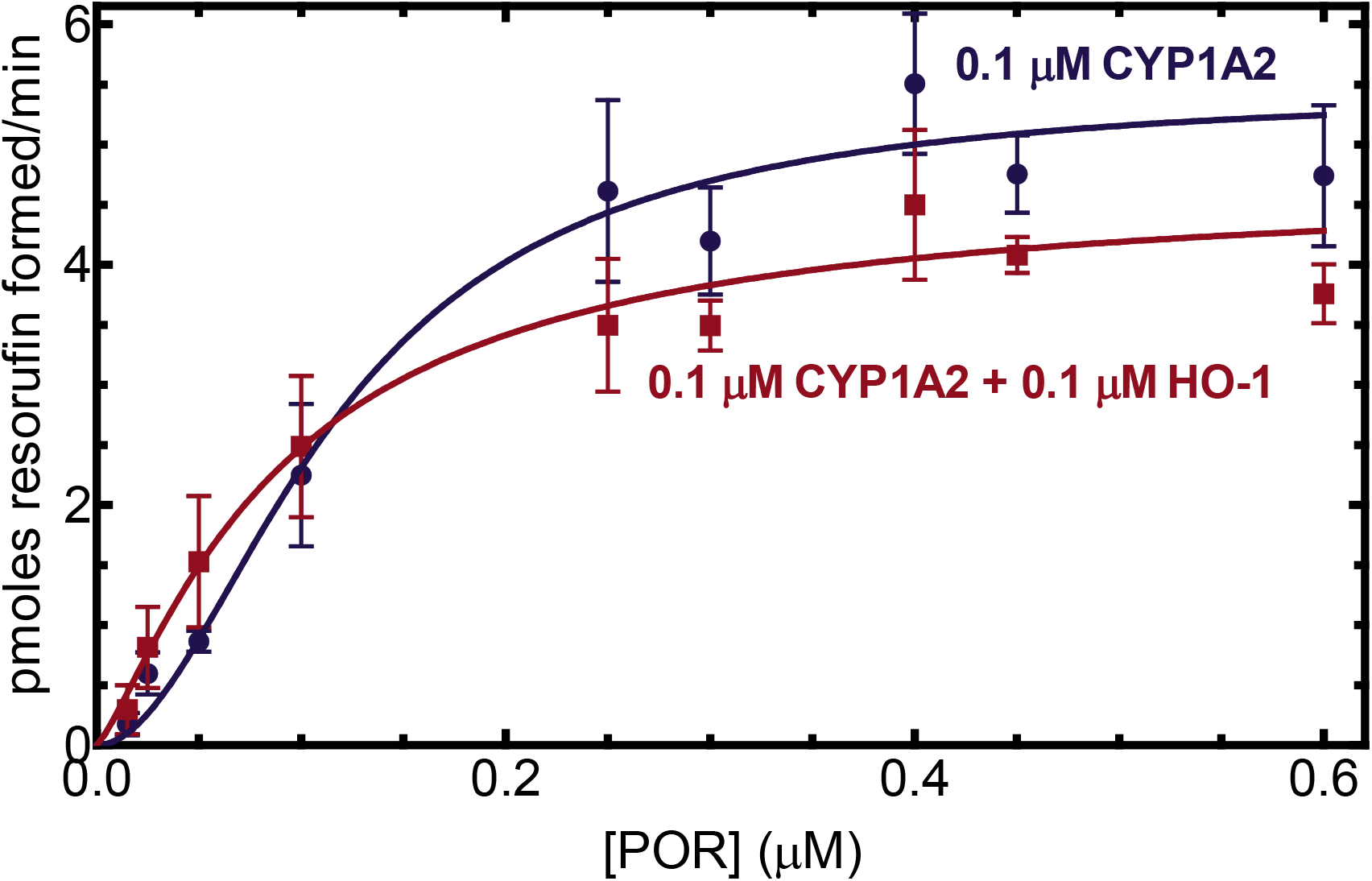
Effect of HO-1 on CYP1A2-mediated 7-ethoxyresorufin O-dealkylation. CYP1A2 (0.1 μM) was reconstituted in DLPC over a range of POR concentrations in the presence and absence of 0.1 μM HO-1. The rates of product formation were determined by measuring fluorescence of the product. Curves were fit to allosteric sigmoidal curves. The curves represent the average of 12 separate experiments to verify the differences observed at both subsaturating and saturating POR. Each data point represents the mean ± SD from at least 4 determinations.

### Effect of CYP1A2 on HO-1 activity

In the converse experiment, the effect of CYP1A2 on HO-1-mediated heme metabolism was examined (Fig. 7). As previously reported (36,37), in the binary reconstituted system containing HO-1 and POR, very tight binding between the proteins was observed having a Km of about 0.001 μM. However, addition of CYP1A2 caused an inhibition of HO-1 activities at subsaturating POR. The apparent K_m_^POR•HO-1^ increased from less than 0.001 μM to 0.024 ± 0.006 μM in the presence of CYP1A2.

**Figure 7.**
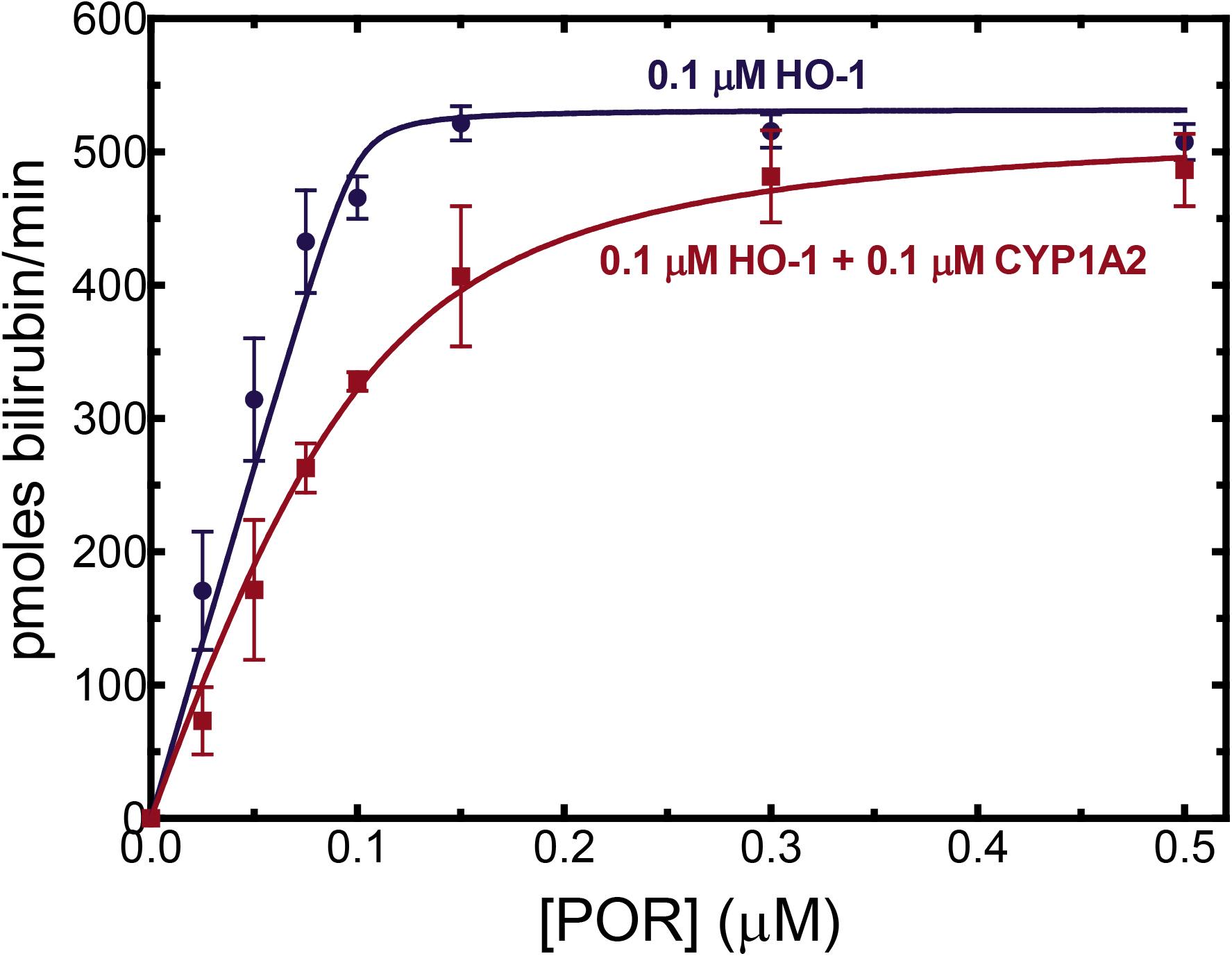
Effect of CYP1A2 on heme degradation by HO-1. HO-1 (0.1 μM) was reconstituted in DLPC at POR concentrations ranging from 0.025-0.5 μM in the absence (green) and presence (brown) of 0.1 μM CYP1A2. Heme oxygenase activity was determined by monitoring the formation of bilirubin using a coupled assay containing biliverdin reductase. The presence of CYP1A2 led to inhibition of HO-1 activity at subsaturating POR but not at higher POR saturation. Data were fit to the Morrison tight binding equation using nonlinear regression. Plotted data represent the mean and SD of 4 determinations.

## Discussion

The P450s and HO-1 share a necessary binding partner in POR. P450s are inducible by a variety of endobiotic and xenobiotic agents (1,2). HO-1 also is inducible by endogenous and foreign compounds as well as by oxidative stress (38). POR is often expressed at much lower levels than are the P450s, leading to conditions where they must compete for limiting POR (39,40); however, the environment can be dramatically changed by HO-1. In the uninduced state, HO-1 levels are lower than those of both POR and total P450; however, when induced, HO-1 concentrations can exceed the levels of both proteins (41). These factors contribute to the potential for P450s, HO-1, and POR to interact differently based on the relative levels of each of the proteins. Thus, different induction states might change how these interactions manifest in terms of enzyme activities.

The simplest interaction model is one where HO-1 and CYP1A2 do not form a complex, and the proteins simply compete for POR based on their relative affinities. We know that HO-1 has an extremely high affinity for POR (K_m_^POR-HO-1^ ~ 0.001 μM) (37,42) & (Fig. 7), whereas the apparent affinity of CYP1A2 for POR is ~60 times lower (K_m_^POR-CYP1A2^ ~ 0.055 μM) (Fig. 6) & (17–19). These data allow us to predict that, when all three proteins are present, HO-1 should outcompete CYP1A2 for the available POR and that CYP1A2 activity should be dramatically inhibited by the addition of equimolar HO-1, which is consistent with the BRET responses observed (Figs. 3 & 4). However, the BRET results also clearly establish that a CYP1A2•HO-1 complex is produced (Fig. 2), eliminating the possibility that these proteins exist in the membrane exclusively as monomers.

A second possibility is that the CYP1A2•HO-1 complex forms but formation of this complex does not change the ability of either moiety to bind with POR. According to this model, an HO-1•CYP1A2 complex would be evident (Fig. 2), the POR/CYP1A2 BRET complex would be significantly disrupted by the presence of unlabeled HO-1, and the high affinity POR•HO-1 complex would be minimally affected by unlabeled CYP1A2. The BRET results (Figs. 3 & 4) are consistent with this mechanism.

The BRET measurements of physical complex formation are consistent with a CYP1A2•HO-1 complex that is capable of forming a high affinity POR•HO-1 complex. This result is corroborated by the significant inhibition of POR•CYP1A2 by HO-1, and the inability of CYP1A2 to disrupt the POR•HO-1 complex. Based on BRET studies alone, a substantial inhibition of CYP1A2-mediated EROD activity would be expected in the presence of HO-1. However, when EROD was measured and POR was limiting, CYP1A2-mediated activity actually increased in the presence of HO-1 despite BRET data showing significant depression of the POR•CYP1A2 complex. CYP1A2-mediated EROD was only inhibited at higher POR concentrations. When CYP1A2 and POR were examined in a simple binary system, CYP1A2 activities exhibited sigmoidal kinetics as a function of the POR concentration (Fig. 6), which was more pronounced at higher [CYP1A2] (12,43). In a previous report, we proposed that CYP1A2 forms a homomeric CYP1A2•CYP1A2 complex that is less active (12). Higher concentrations of POR disrupt this complex, and allow the formation of a functional POR•CYP1A2 complex. In like manner, the binding of HO-1 may disrupt the inhibitory CYP1A2•CYP1A2 complex, allowing generation of functional POR•CYP1A2 at low POR concentrations. Although part of the increased EROD could be rationalized by the disruption of an inhibitory CYP1A2•CYP1A2 complex, it does not reconcile with the dramatic disruption of the POR•CYP1A2 complex by HO-1 (Fig. 3). In other words, if HO-1 is present and disrupts the CYP1A2•CYP1A2 complex, it would be replaced by the CYP1A2•HO-1 complex, where the HO-1 moiety would preferentially bind POR. At higher POR concentrations, inhibition of EROD was observed, suggestive of a conformationally mediated alteration in the maximum rate of EROD catalysis upon formation of the CYP1A2•HO-1 complex.

CYP1A2 also was shown to affect HO-1 mediated heme metabolism (Fig. 7). Based on the BRET studies alone, HO-1 activities were expected to be negligibly affected by the presence of CYP1A2, particularly at subsaturating POR (Fig. 4). Although the maximal response was not affected at excess POR concentrations, HO-1-mediated catalysis was inhibited by CYP1A2 at subsaturating POR. This inhibition of HO-1 mediated heme metabolism at subsaturating POR would be unexpected unless formation of the CYP1A2•HO-1 complex conformationally affects the catalytic characteristics of the component enzymes.

These results provide important information regarding the interactions between the HO-1 and P450 systems, but also raise many important questions. First, complex formation between HO-1 and different P450s is specific; not all P450s appear to form such complexes (Fig. 1). Second, the BRET response using CYP1A2-GFP and Rluc-HO-1 shows the expected saturation as the ratio of the GFP-to Rluc-containing proteins are increased (Fig. 2). Interestingly, this complex is not significantly affected by the addition of POR, which indicates that, when both proteins are present, they act not as monomers or homomeric complexes, but as a ternary (or higher order) complex. Third, the functional POR•CYP1A2 BRET complex is affected by HO-1; however, the POR•HO-1 complex is not affected by CYP1A2. Finally, the activities of both CYP1A2-mediated EROD and HO-1-mediated heme metabolism are influenced by the presence of the other protein in ways that cannot be explained by a simple mass-action competition for limiting POR. The data at this point suggest multiple effects on the oligomerization/conformation of the enzymes in the ternary systems. A more detailed kinetic analysis will be required to identify the specific modifications taking place.

## Experimental Procedures

### Materials

Dulbecco’s Modified Eagle’s Medium (DMEM), phosphate buffered saline (PBS), fetal bovine serum (FBS), and Lipofectamine 2000 were purchased from Invitrogen (Eugene, OR). The plasmids used to generate BRET vectors (pGFP^2^-N1, pGFP^2^-N2, pRluc-N2, pRluc-N3) and the pGFP^2^-Rluc vector were obtained from BioSignal Packard (Waltham, WA). GFP^2^ is a wild-type green fluorescent protein (GFP) that has been modified by a F64L substitution mutation which results in brighter fluorescence but similar excitation and emission spectra. Human embryonic kidney (HEK)-293T/17 cells were obtained from ATCC (Manassas, VA). Antibiotic-antimycotic solution was purchased from Life Technologies (Carlsbad, CA). Coelenterazine 400A was purchased from Gold Biotechnology (St. Louis, MO), and coelenterazine h was purchased from Promega (Madison, WI). 7-ethoxyresorufin was purchased from Anaspec (Fremont, CA), and cytochrome c was obtained from Sigma (St. Louis, MO).

### Generation of BRET vectors

HO-1, rabbit CYP1A2, and POR were wild-type proteins without any modifications to their amino acid sequences. To generate the BRET expression vectors, linear cDNA coding for full-length, wild-type CYP1A2 (NM_001171121), and POR (NM_001367562) was amplified from existing bacterial expression vectors with PCR using primers to introduce restriction sites that immediately flanked the full-length gene, excluding the stop codon. The restriction sites (with the 5’ site listed first) were: EcoRI and BamHI for CYP1A2, and EcoRI and HindIII for POR. These PCR products were then ligated into the empty vectors (pGFP^2^-N1 and pRluc-N2 for the P450s, pGFP^2^-N3 and pRluc-N1 for POR) after each was digested by the indicated pair of restriction enzymes (New England Biolabs, Inc.; Ipswich, MA). The multiple cloning sites of the vectors are upstream of the GFP or Rluc tag. This information is summarized in Table 1. This orientation was chosen to ensure that the heterologous protein tags were less likely to interfere with the ability of the membrane-binding regions of the proteins to insert into the ER membrane. Due to the restriction sites used, the BRET vectors code for a 5-20 amino acid sequence between the 3’ end of the inserted gene and the start codon for the GFP or Rluc tag. To generate vectors for the expression of untagged, wild-type CYP1A2, HO-1, and POR, site-directed mutagenesis was performed to create a stop codon immediately 3’ to the main protein sequence. Site-directed mutagenesis was performed using the QuikChange II kit from Agilent Technologies (Santa Clara, CA). All PCR amplification and mutagenesis primers were purchased from Integrated DNA Technologies (Coralville, IA). Insertion of cDNA and mutagenesis were confirmed by sequencing (ACGT; Germantown, MD).

**Table 1 –.** Description of the BRET constructs and their restriction sites. Each of the GFP- and Rluc-labeled P450, HO-1 and POR constructs were cloned into the indicated parent vectors using the restriction site shown. The BRET vectors were purchased from BioSignal Packard (Waltham, MA).

| Construct | Parent vector | 5' site | 3' site | Sequence identifier | Peptide linker |
| --- | --- | --- | --- | --- | --- |
| CYP1A1-GFP | pGFP<sup>2</sup>-N3 | MluI | SacII | BC023019 | PRARDPPVAT |
| CYP2D6-GFP | pGFP<sup>2</sup>-N3 | MluI | SacII | BC067432.1 | PRARDPPVAT |
| CYP2A6-GFP | pGFP<sup>2</sup>-N3 | MluI | SacII | BC067430.1 | PRARDPPVAT |
| CYP2C9-GFP | pGFP<sup>2</sup>-N3 | MluI | SacII | BC125054.1 | PRARDPPVAT |
| GFP-HO-1 | pGFP<sup>2</sup>-C3 | SacI | XhoI | NM_002133.3 | SGLRSGEL |
| Rluc-HO-1 | pRluc-C3 | SacI | XhoI | NM_002133.3 | RARDPGLRSGEL |
| Unlabeled HO-1 | pGFP<sup>2</sup>-N1 | SacI | BamHI | NM_002133.3 | N/A |
| POR-GFP | pGFP<sup>2</sup>-N2 | EcoRI | BamHI | NM_001367562 | GSPPVAT |
| POR-Rluc | pRluc-N3 | EcoRI | BamHI | NM_001367562 | GSTGAT |
| Unlabeled POR | POR-GFP | EcoRI | BamHI | NM_001367562 | N/A |
| CYP1A2-GFP | pGFP<sup>2</sup>-N1 | EcoRI | BamHI | NM_001171121.1 | WIPPVAT |
| CYP1A2-Rluc | pRluc-N2 | EcoRI | BamHI | NM_001171121.1 | WIPTGAT |
| Unlabeled CYP1A2 | CYP1A2-GFP | EcoRI | BamHI | NM_001171121.1 | N/A |

Vectors coding for human CYP1A1 (vector: pCMV-SPORT6; accession number: BC023019), CYP2A6 (pCR-BluntII-TOPO; BC067430.1), CYP2C9 (pCR-BluntII-TOPO; BC125054.1), and CYP2D6 (pDNR-Dual; BC067432.1) were purchased from GE Healthcare/Dharmacon (Lafayette, CO). The pPORhWT vector that served as a source of human POR cDNA was a generous gift of Dr. Bettie Sue Masters (44). Human HO-1 cDNA was derived from the pGEX-4T-2 vector described previously (37).

Because the C-terminus of HO-1 binds to cellular membranes (in contrast with the P450s and POR) (45,46), N-terminal tags were used to avoid interfering with membrane insertion. This involved using the pRluc-C3 and pGFP^2^-C2 vectors with 5’ SacI and 3’ XhoI restriction sites and designing primers that conserved the 3’ stop codon. In order to create a vector for unlabeled HO-1 expression, the coding sequence was excised from the pRluc-C3 vector again using the 5’ SacI site with a 3’ BamHI site downstream of the stop codon. The complete vector was then produced by ligating the HO-1 insert into a similarly digested pGFP^2^-N1 backbone.

### BRET assays

For BRET assays, cells were first transfected with 1-3 μg total DNA at different ratios of the GFP and Rluc constructs. The goal for these transfections was to generate a series of cells expressing the same total protein at a range of GFP:Rluc ratios to ensure that the BRET complexes were specific and not due to changes in protein expression levels. If this transfection strategy yielded approximately the same total protein expression across all conditions, the results were considered valid. In many cases, however, one or two additional trials were necessary to achieve constant levels of protein expression. After allowing at least 24 hours for protein expression, cells were checked using fluorescence microscopy to ensure efficient expression of the GFP fusion proteins. Each transfection was performed with the following three controls: a GFP-Rluc fusion protein, the Rluc fusion protein alone, and cells transfected with only pUC19. Cells were harvested in 1 ml PBS, centrifuged, and resuspended in 700 μl PBS. For BRET measurements, 100 μl of suspended cells were distributed in quadruplicate into an opaque, white 96-well plate (PerkinElmer; Waltam, MA). A TriStar LB 941 microplate reader (Berthold Technologies; Bad Wildbad, Germany) was used for BRET measurements. This plate reader was programmed to perform the following actions for each well: inject 100 μl of a 10 μM coelenterazine 400A/PBS solution, shake for 1s to mix, read Rluc emission for 3s at 410 nm, and read GFP emission for 3s at 515 nm.

### Calculation of the net BRET ratio

The ratio of GFP fluorescence (at 510 nm) to Rluc luminescence (at 410 nm) immediately following the addition of coelenterazine 400a to a final concentration of 5 μM was used as the BRET measurement. For each transfection condition, net GFP and Rluc emissions were calculated by comparing with the respective signals of untransfected cells. The raw BRET ratio was then compared to the BRET signal from cells expressing only the Rluc-tagged protein to obtain BRETnet (10,12).

### Determination of relative protein expression

Relative expression levels of GFP and Rluc tagged proteins were determined photometrically by comparison to cells expressing a GFP-Rluc fusion protein. First, GFP expression was determined by measuring fluorescence (410 nm excitation; 515 nm emission) on a SpectraMax M5 plate reader using 100 μl samples of the original cell suspension in black clear-bottomed 96-well plates (Corning Inc.; Corning, NY). Rluc expression was measured in the microplate reader using the fourth quadruplicate of each experimental condition. Expression of the Rluc fusion protein was estimated by the addition of coelenterazine h to a final concentration of 5 μM, and unfiltered emission was measured for 1 s. The GFP and Rluc signals for each sample were then normalized to that of the GFP-Rluc fusion protein (which is assumed to have a 1:1 GFP:Rluc expression ratio), so that dividing the normalized GFP value by the normalized Rluc value yielded an approximation of the actual GFP:Rluc expression ratio. In each case, there were no significant differences when comparing the data in the absence and presence of the unlabeled protein.

### Preparation of reconstituted systems for enzyme activity measurements

Reconstituted systems were prepared as described previously (36). A 5 mM DLPC stock solution was prepared in 50 mM potassium phosphate (pH 7.25) with 20% (v/v) glycerol, 0.1 M NaCl, and 5 mM EDTA. This solution was then sonicated at room temperature in a water-bath sonicator for 20 minutes or until the solution clarified. To set up POR titration assays, purified recombinant CYP1A2, HO-1, or both were mixed with varying quantities of purified recombinant POR and preincubated for 2 h at room temperature in DLPC stock solution. During preincubation, enzyme concentration was kept as high as possible and (unless otherwise noted) the molar ratio of DLPC to the enzyme being assayed was kept at 160:1 (19).

### HO-1 activity assay

To measure HO-1 activity, we used a coupled assay including a source of biliverdin reductase, which converts the biliverdin generated by HO-1-mediated heme catabolism into bilirubin. Reconstituted systems (RS) were prepared and pre-incubated for 2 h at room temperature before the addition of other reaction components in 0.1 M potassium phosphate (pH 7.4). The substrate, heme, was added at 15 μM and rabbit liver cytosol (0.8 mg/ml protein) to provide biliverdin reductase. Catalase was added to a concentration of 0.25 U/μl in order to mitigate the effect of H_2_O_2_ accumulation on reaction linearity (36). All activity assays were allowed to pre-incubate at 37°C for 3 min prior to initiation of the reaction by the addition of NADPH to a final concentration of 0.5 mM. Each reaction was performed at 37°C, in triplicate, in black, clear-bottom 96-well plates with a final volume of 0.1 ml. A SpectraMax M5 microplate reader was used to monitor bilirubin formation using a real-time assay, measuring the difference in absorbance at 464 and 530 nm. The assay was monitored for 10 min and the initial rates measured from the slope from the linear portion of the curve. Enzyme activities were calculated using the delta extinction coefficient of 40 mM^-1^(cm)^-1^ (47,48).

### 7-Ethoxyresorufin-O-deethylase assay

For the measurement of CYP1A2 activity in reconstituted systems, we used 7-ethoxyresorufin (7-ER) as a substrate. CYP1A2 converts 7-ER into the fluorescent product resorufin. After preincubation for 2 h at room temperature, reconstituted systems were diluted into buffer A (50 mM HEPES (pH 7.5), 15 mM MgCl_2_, 0.1 mM EDTA). 7-ER was added to a final concentration of 4 μM. Once at 37 °C, the reactions were initiated with the addition of NADPH to a final concentration of 0.5 mM. Reactions were carried out in real-time at 37°C using a SpectraMax M5 plate reader to monitor resorufin fluorescence (excitation: 535 nm, emission 585 nm). Initial rates were used for calculation of activities using a standard curve generated from known quantities of resorufin.

### Data analysis

Unless otherwise noted, POR titration data were fit to the Morrison equation for tight binding in GraphPad Prism 5 using nonlinear regression (49). Statistical comparison of each estimated parameter was done using an extra-sum-of-squares F test between a simple model in which all conditions share one value for the given parameter and another model, which allows each condition a separate value for the parameter (50). Parameters were considered significantly different if the model with separate values was preferred at p < 0.05. Data points are represented as the mean ± SD.

## Data Availability

All data are contained within the manuscript.

## Acknowledgements

We would like to thank Lucy Waskell (Univ. Michigan, Ann Arbor, MI) for the NADPH-cytochrome P450 reductase expression system, and Mahin Maines (University of Rochester, Rochester, NY) for the HO-1 cDNA. We also would like to thank Marilyn Eyer for technical support with protein purification. A portion of this study was published in a dissertation by Dr. J. Patrick Connick entitled, <u>The cytochromes P450, heme oxygenase-1, and NADPH-cytochrome P450 reductase form multiple complexes that influence protein function”</u>.

## Funding and additional information

This work was supported by NIH grants from the National Institute of General Medical Sciences (R01 GM123253, and the National Institute of Environmental Health Sciences (P42 ES013648). We also would like to thank the Board of Regents of the State of Louisiana in partial support of JPC.

